# Human medial temporal lobe neurons link future reward coding with intertemporal choice and impulsivity

**DOI:** 10.64898/2026.07.09.737293

**Authors:** Marcel S. Kehl, Stefan Dürschmid, Valeri Borger, Rainer Surges, Florian Mormann

## Abstract

The ability to delay gratification emerges early in life and is linked to long-term health and economic success. Conversely, high impulsivity, marked by a preference for immediate rewards, can be associated with psychiatric disorders. Although processes underlying human delay discounting have been studied at behavioural and macroscopic neural levels, they remain elusive at the single-neuron level. Specifically, it remains unknown how human neurons encode extended reward delays, predict intertemporal choices, and how these processes vary with impulsivity. Here, we record single-neuron activity in the human medial temporal lobe (MTL) to explore decision and delay coding. We identify neurons that predict upcoming decisions in the amygdala and hippocampus. Neurons in the entorhinal cortex and hippocampus encode reward delays, with particularly hippocampal population activity coding prospective temporal periods. Importantly, neuronal activity in impulsive individuals shows diminished prospective temporal coding and predicts decisions only shortly before choices are reported. Our findings reveal how distinct MTL regions contribute to intertemporal decisions and provide insight into the neuronal signatures underlying impulsivity.

## Introduction

Humans regularly face decisions between immediate and delayed gratification, such as choosing to watch a movie or prepare for tomorrow’s meeting. Impulsive individuals tend to prefer smaller-but-sooner over larger-but-later rewards, a behaviour termed delay discounting in intertemporal decision-making ^1^. This tendency toward impulsive decision-making can, in severe cases, contribute to psychiatric disorders like addiction ^2,3^. Understanding the neuronal mechanisms underlying impulsivity is essential for advancing our understanding of these disorders ^4,5^.

Non-invasive bulk-imaging techniques have identified the human hippocampus and amygdala as contributors to intertemporal choices ^6,7^. However, the specific mechanisms and roles of different human medial temporal lobe regions in these choices remain unclear at a mechanistic, neuronal level. The amygdala is essential for emotional processes ^8–11^ and plays an important role in decision-making ^12–17^, reward anticipation and the ability to delay gratification ^18^. For instance, individuals with bilateral amygdala damage exhibit reduced aversion to monetary loss ^13^, and recordings from nonhuman primates have demonstrated that amygdala neurons contribute to economic decision-making ^16,17^. The hippocampus, on the other hand, is integral for episodic memory ^19–23^ and also contributes to human decision-making ^24–26^. The hippocampus appears particularly important when intertemporal decisions rely on episodic future thinking, as patients with hippocampal damage can show normal delay discounting ^27,28^, but do not benefit from imagining personally relevant future events, which reduces discounting in healthy individuals ^25,29^. The mechanisms underlying future episodic thinking and prospective temporal coding remain largely unknown at the neuronal level ^7,28^. Specifically, it is unclear how reward-delay information and individual decisions are encoded by human MTL neurons.

A key aspect of episodic future thinking is representing the time until future events occur (e.g., three months from now). MTL neurons in rodents ^30^ and humans ^31^ encode time intervals spanning seconds to minutes (‘time cells’). Whether and how prospective temporal coding differs between impulsive and non-impulsive individuals remains an open question. Identifying such differences could reveal how MTL reward-delay representations support future planning and how their disruption contributes to impulsive choice.

Using the rare opportunity to record single-neuron activity in neurosurgical patients, we investigated how individual neurons in the human MTL contribute to intertemporal choices. We find that the human MTL contains decision-coding and delay-coding neurons and that neuronal population activity encodes both reward delays and trial-by-trial decisions. Region-specific decoding shows that reward delays are most reliably encoded by hippocampal neurons, while both amygdala and hippocampal neurons predict upcoming decisions. Notably, prospective temporal coding is observed only in non-impulsive individuals. Impulsive individuals show decision coding only shortly before decisions are reported, whereas non-impulsive individuals demonstrate earlier and more sustained decision coding. Our results provide insights into the neuronal coding mechanisms during intertemporal choices across MTL regions and the neural basis of impulsive behaviour.

## Results

### Human single-neuron recordings during intertemporal choices

To explore the mechanisms underlying intertemporal decision-making within the human MTL at the neuronal level, we recorded the activity of individual neurons in 17 epilepsy patients undergoing invasive seizure monitoring during a delay-discounting task (Figure 1A & B). Overall, we recorded the activity of 1,002 neurons in the human MTL (Figure 1C & D). Electrodes were implanted bilaterally in the amygdala (Am), hippocampus (Hp), entorhinal cortex (EC) and parahippocampal cortex (PHC) (Figure 1C & Figure S2). During the delay discounting task, participants were asked to choose between a smaller-but-sooner (SS) reward and a larger-but-later (LL) reward of 10 € at variable delays (1, 2, 5, 11, 24 or 52 weeks in a total of 90 trials) (Figure 1E). Using an adaptive delay-discounting paradigm, we estimated each participant’s discounting behaviour and observed a broad range from high to low impulsivity, quantified by the discounting parameter k (Methods; Figure 1F).

**Figure 1.**
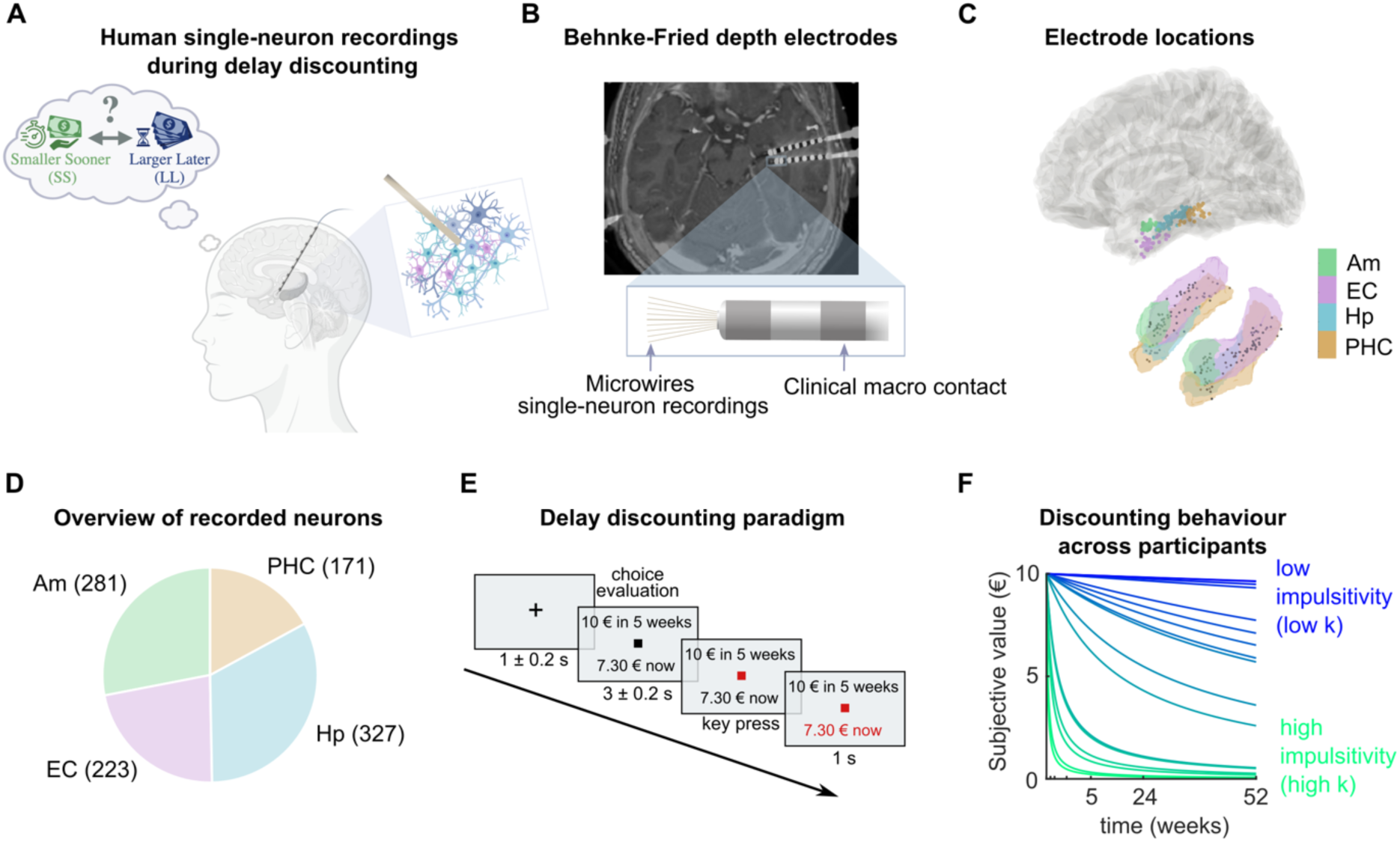
Recording human single-neuron activity during intertemporal decisions. (A) Illustration of single-neuron recordings in humans during intertemporal choices. Participants were asked to decide between a larger-but-later and a smaller-but-sooner reward while neuronal activity within the MTL was recorded. Created in BioRender. Kehl, M. S. (2026) https://BioRender.com/opg4ex1 (B) Top: Co-registered MRI and CT scans with implanted Behnke-Fried depth electrode in a neurosurgical patient. Bottom: Illustration of a Behnke-Fried depth electrode showing the two innermost clinical macro contacts as well as microwires protruding from the tip of the electrode. Am, amygdala; EC, entorhinal cortex; Hp, hippocampus; PHC, parahippocampal cortex. (C) Top: Electrode locations across all 17 patients. Coloured dots indicate the positions of innermost clinical macro contacts projected onto the Montreal Neurological Institute (MNI) ICBM152 template ^32^. Bottom: Electrode locations projected onto a 3D reconstruction of the targeted MTL regions based on the Brainnetome Atlas ^33^. (D) Number of neurons recorded for each anatomical target region, coloured as in (C). (E) Delay discounting paradigm: Participants decided between an SS (here: 7.30 €) and an LL reward (10 € at 6 different delays) across 90 trials. The discounting parameter k, which determines the equivalent SS amount based on the delay, was dynamically adjusted according to previous choices to assess individual discounting. (F) Discounting behaviour across all 17 participants. For each participant (coloured lines) the hyperbolic discounting was estimated based on the individual discounting parameter k. High impulsivity (green) is indicated by strong discounting of future rewards, while low impulsivity (blue) is marked by more stable subjective value across delay.

### Human amygdala and hippocampal neurons predict intertemporal decisions

The human MTL is involved in intertemporal choices ^6,7^. However, whether and how individual neurons in the human MTL encode and contribute to intertemporal decisions remains poorly understood. We therefore asked whether the firing of human MTL neurons predicts intertemporal decisions during the choice-evaluation interval before decisions were reported. Comparison of the firing rates of MTL neurons during trials in which SS vs. LL options were chosen revealed that a significant fraction of MTL neurons adjusted their firing based on individuals’ decisions (Figure 2A, 77 out of 1,002 neurons, P = 0.00017, right-sided binomial test). These decision-coding neurons were specifically observed in the amygdala and hippocampus, but not in the EC and PHC (Figure 2A, Am: 8.2%, 23 of 281, P = 0.015; Hp: 9.2%, 30 of 327, P = 0.0012; EC: 5.4%, 12 of 223, P = 0.44; PHC: 7%, 12 of 171, P = 0.15, right-sided binomial test). Figure 2B depicts an amygdala neuron that increased firing during the choice-evaluation interval, when the SS option was chosen. Figure 2C & D show two hippocampal neurons, one exhibiting increased activity for the LL reward (Figure 2C), whereas the other was more active when the SS reward was chosen (Figure 2D). Individual neurons modulated their firing in relation to upcoming decisions and displayed rich, heterogeneous response profiles, suggesting that decision information emerges from distributed and coordinated activity across the neuronal population. We therefore tested whether this population-level activity could predict upcoming choices by performing a neuronal decoding analysis based on the spiking activity of the recorded MTL neurons ^34^, which revealed significant decision coding while participants evaluated choice options (Figure 2E). The decision was decodable across a sustained interval, starting around 1 s after the onset of the choice-evaluation window. Region-specific decoding demonstrated that choices were most reliably predicted by neurons in the human amygdala and hippocampus (Figure 2F). Notably, amygdala neurons predicted upcoming decisions already in the time interval before choice options were presented (Figure 2F).

**Figure 2.**
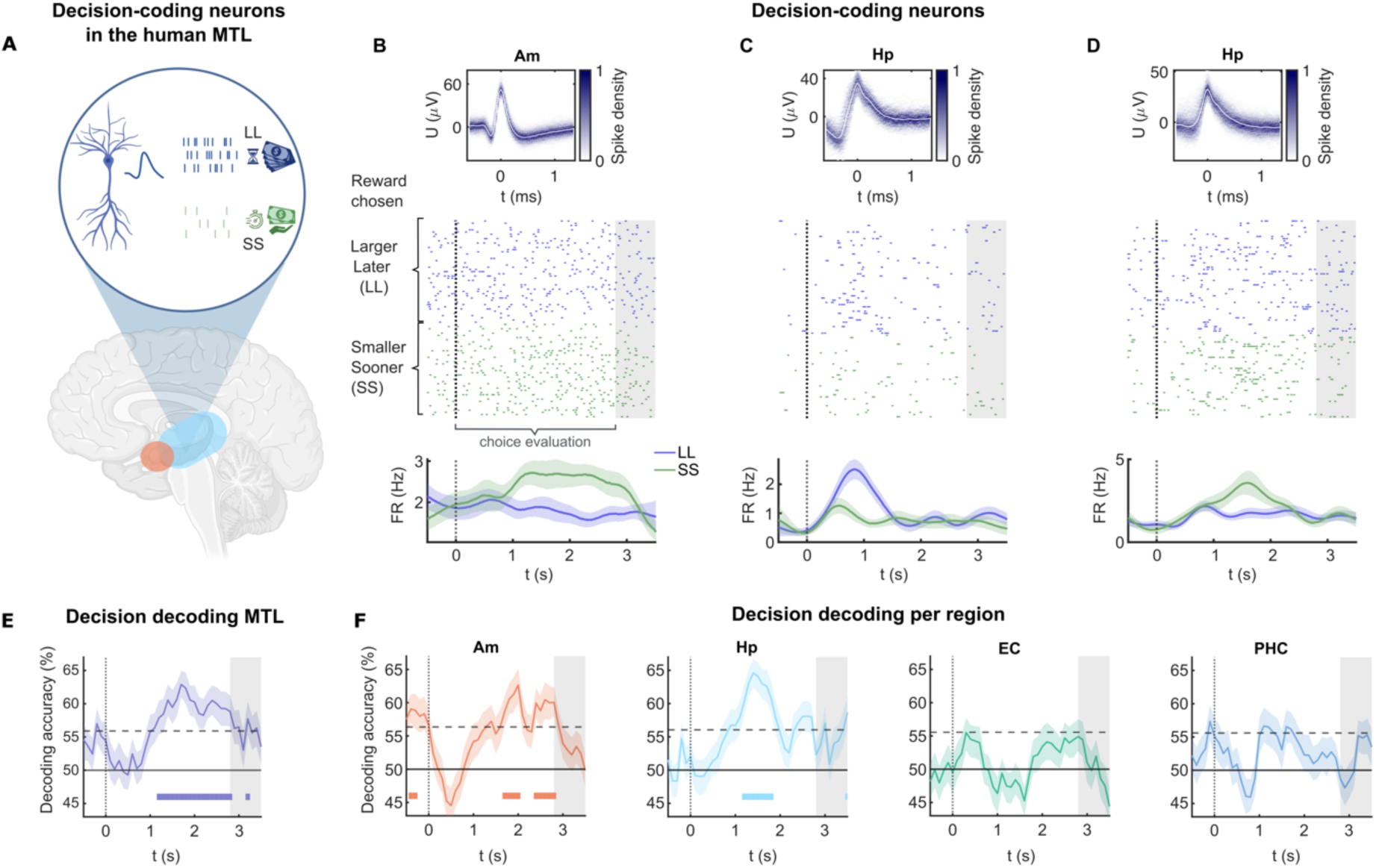
Human amygdala and hippocampal neurons predict intertemporal decisions. (A) Illustration of decision coding in a human MTL neuron. Decision-coding neurons change their firing rate (FR) during the choice-evaluation interval depending on whether the SS or LL option was chosen. A significant fraction of decision-coding neurons was recorded in the human MTL (77 out of 1,002 neurons, P = 0.00017, right-sided binomial test). A significant fraction of decision-coding neurons was observed in the human amygdala and hippocampus (Am: 8.2%, 23 of 281, P = 0.015; Hp: 9.2%, 30 of 327, P = 0.0012; EC: 5.4%, 12 of 223, P = 0.44; PHC: 7%, 12 of 171, P = 0.15, right-sided binomial test). Created in BioRender. Kehl, M. S. (2026) https://BioRender.com/4u0er67 (B-D) Examples of decision-coding neurons in the human amygdala and hippocampus. Top: Neuronal spike-shape density (mean ± s.d. in white). Middle: Raster plot locked to choice-option onset (t = 0 s) with the grey shaded area marking the choice-reporting period (beginning at 2.8 – 3.2 s). Bottom: Averaged firing rate (mean ± s.e.m., 1 s Gaussian kernel smoothing) for trials in which the LL (blue) or SS (green) options were chosen. (B) Amygdala neuron that increased its firing in trials in which the SS option was chosen (two-sided t-test, P = 0.003, t(88) = 3.1). (C) Hippocampal neuron that increased its firing during trials in which the LL option was chosen (two-sided t-test, P = 0.015, t(88) = -2.5). (D) Hippocampal neuron with increased firing during SS trials (two-sided t-test, P = 0.014, t(88) = 2.5). (E) The population activity of human MTL neurons predicted intertemporal decisions during choice evaluation. Decoding accuracy (mean ± bootstrapped 95% confidence interval (CI) across resampling runs) is shown in blue with chance level performance (50%) indicated by the solid horizontal line. Decoders were evaluated using 10 cross-validation data splits and 100 resampling runs each using 1 s time bins and a 100 ms step size. Times of significant decision coding were determined based on label-shuffled data (1,000 label-permutations for each of the 41 time bins), with the dashed horizontal line indicating the 95^th^ percentile of label-permuted decoding performances across all 41,000 decoders. Thick horizontal bars indicate times of significant decoding based on the percentile of the real decoding results within the label-permuted distribution and after correcting for multiple comparisons across all 41 time bins using false-discovery-rate (FDR) correction. (F) Decision coding for neurons across anatomical target regions (decoding and display as in E). Significant decision coding after FDR correction was observed in the human amygdala and hippocampus (thick horizontal bars). Amygdala neurons exhibited significant decision coding already before presentation of the choice options.

Together, our recordings reveal that intertemporal decisions are predicted by the activity of individual neurons in the human MTL, specifically in the amygdala and hippocampus, with amygdala neurons showing anticipatory coding of decisions.

### Prospective temporal coding of reward delays in the human MTL

The human MTL is critical for future planning ^25,29,35,36^ and the representation of temporal information ^37^. The mechanisms of temporal coding across extended future time periods are not well understood at the neuronal level in humans. To address this, we examined whether human MTL neurons encode temporal information of choice options (i.e., reward delays) across extended prospective time intervals of up to one year.

We found that a significant fraction of human MTL neurons modulate their firing rates based on the delay of future rewards (Figure 3A, 63 neurons, P = 0.04, right-sided binomial test). These reward-delay coding neurons were most frequently found in the hippocampus and EC (Figure 3A, Am: 4.6%, 13 of 281, P = 0.65; Hp: 7.3%, 24 of 327, P = 0.041; EC: 8.5%, 19 of 223, P = 0.017; PHC: 4.1%, 7 of 171, P = 0.76, right-sided binomial test). To further explore MTL delay coding, we asked whether MTL neurons’ firing rates are correlated with delay duration and found that a significant fraction of human MTL neurons exhibited such a correlation (Figure 3A, 74 neurons, P = 0.00068, right-sided binomial test). Such delay-correlated neurons were specifically found in the hippocampus (Figure 3A, Am: 7.1%, 20 of 281, P = 0.073; EC: 6.3%, 14 of 223, P = 0.23; Hp: 9.2%, 30 of 327, P = 0.0012; PHC: 5.8%, 10 of 171, P = 0.35). Figure 3B shows a hippocampal neuron that exhibits increased firing for longer reward delays, whereas another hippocampal neuron responded more strongly for short delays (Figure 3C). Figure 3D displays an EC neuron that does not exhibit a monotonic relationship with delay duration. Instead, it shows a U-shaped response, firing most strongly for the shortest and longest reward delays.

**Figure 3.**
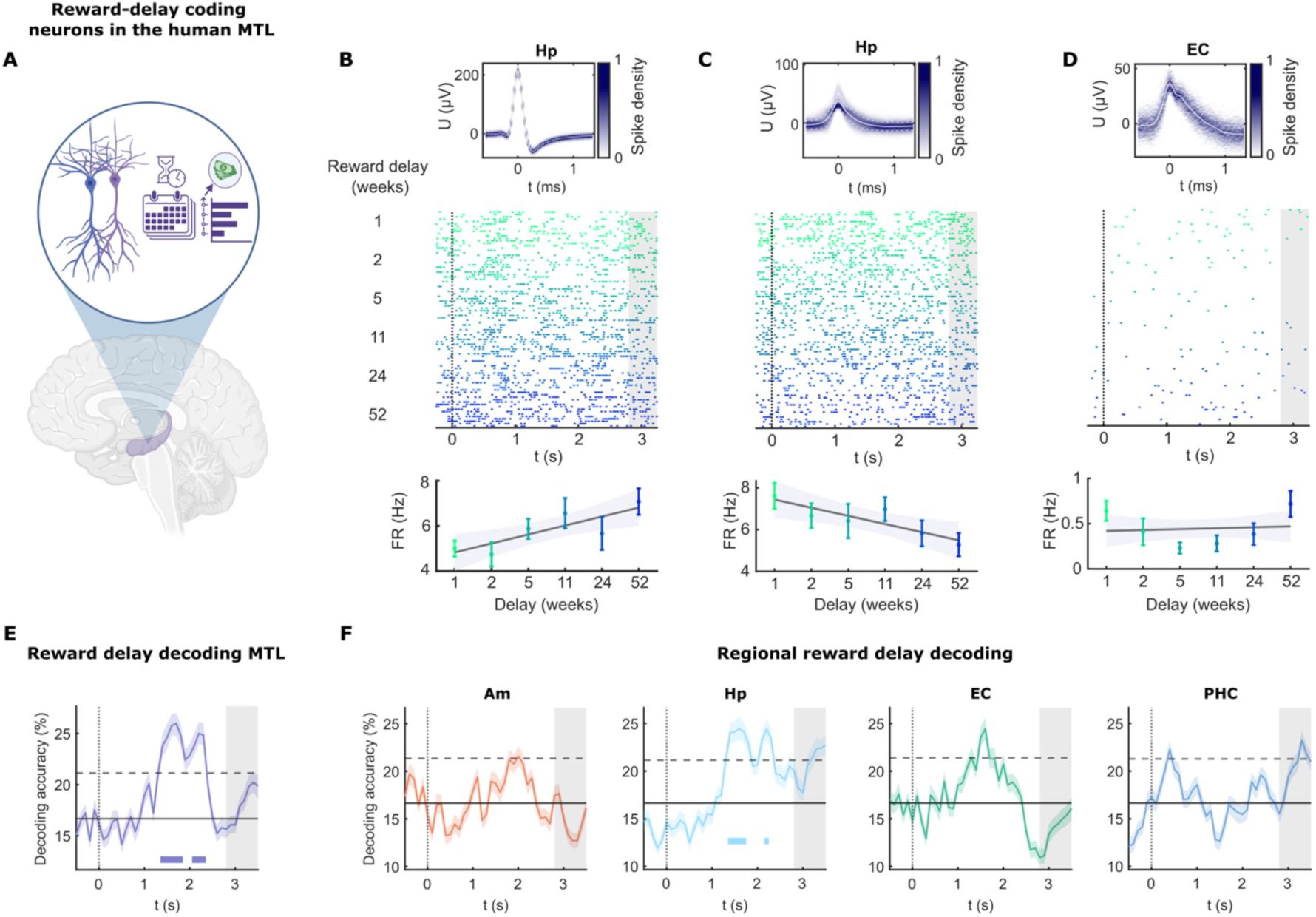
Prospective temporal coding of reward delays in the human MTL. (A) MTL neurons modulate their firing rate based on future reward delays. Overall, 63 reward-delay coding neurons were recorded in the MTL (P = 0.04, right-sided binomial test). They were most frequently found in the hippocampus and EC (Am: 4.6%, 13 of 281, P = 0.65; Hp: 7.3%, 24 of 327, P = 0.041; EC: 8.5%, 19 of 223, P = 0.017; PHC: 4.1%, 7 of 171, P = 0.76, right-sided binomial test). In addition, 74 delay-correlated neurons showed a significant Spearman correlation with reward-delay duration (P = 0.00068, right-sided binomial test), primarily in the hippocampus (Am: 7.1%, 20 of 281, P = 0.073; EC: 6.3%, 14 of 223, P = 0.23; Hp: 9.2%, 30 of 327, P = 0.0012; PHC: 5.8%, 10 of 171, P = 0.35). Created in BioRender. Kehl, M. S. (2026) https://BioRender.com/tkx0u1h (B-D) Top: Neuronal spike-shape density (mean ± s.d., white). Middle: Raster plot locked to choice-option onset (t = 0 s) with the grey shaded area marking the choice-reporting period (beginning at 2.8 – 3.2 s). Reward delays colour-coded from top to bottom (1 week up to 52 weeks, green to blue). Bottom: Firing rate per delay (mean ± s.e.m.). Linear regressions (grey) with 95% confidence intervals (light blue). (B) Hippocampal neuron that changes its firing rate based on reward delay, responding more strongly to larger delays (one-way ANOVA of spike count with reward delay, P = 0.038, F(5,84) = 2.5; Spearman correlation ρ = 0.28, P = 0.0074, N = 90). (C) Hippocampal neuron that increased its firing for short reward delays (Spearman correlation ñ = -0.23, P = 0.0281, N = 90; one-way ANOVA, P = 0.14, F(5,84) = 1.7). (D) Example of a reward-delay coding neuron in EC that shows no monotonic relation between firing rate and delay but exhibits increased activity for both long and short reward delays (Spearman correlation ñ = 0.008, P = 0.94, N = 90, one-way ANOVA, P = 0.021, F(5,84) = 2.8). (E) Reward-delay coding in human MTL neurons: During the evaluation interval (starting at t = 0 s, dotted vertical line, until t = 2.8 – 3.2 s) both choice options are evaluated by participants. Decisions are indicated during the reporting interval (grey shaded area). Decoding accuracy (mean ± bootstrapped 95% CI across resampling runs, 1 s time bins with 100 ms step size) is shown in blue with chance level performance (16.67%, i.e., 1 out of 6 delays) indicated by the solid horizontal line. Decoders were evaluated using 10 cross-validation data splits and 100 resampling runs each. Significant delay coding is observed during the evaluation interval. Times of significant delay coding were determined based on label-permuted data (1,000 label-permutations for each of the 41 time bins), with the dashed horizontal line indicating the 95^th^ percentile of label-permuted decoding performances across all 41,000 decoders. Thick horizontal bars indicate times of significant decoding based on the percentile of the real decoding results within the label-permuted distribution and after correcting for multiple comparisons across all 41 time bins using FDR correction. (F) Reward-delay coding for neurons across anatomical target regions. Display and decoding as in (E). The highest delay coding was observed in the hippocampus and EC, with significant delay coding in the hippocampus after FDR correction.

We next asked how the MTL population activity encodes reward-delay information. Population decoding based on the spiking of the recorded MTL neurons demonstrated significant temporal coding during the evaluation of choice options (Figure 3E). Future reward delays could be decoded above chance about 1.5 seconds after both options appeared on the screen, allowing prior evaluation of both options ^38^. The overall delay decoding was not dominated by any single delay. Instead, both long and short delays contributed to the decoding performance (Figure S6). Region-specific decoding revealed that reward delays were most accurately predicted by hippocampal neurons (Figure 3F). Combined, these results demonstrate that human MTL neurons represent the temporal structure of future reward delays across extended periods.

To further explore how delay and decision information are integrated, we applied a generalized mixed-effects model to predict neuronal activity based on delay, decision, and their interaction (Figure S7, Table S2). This analysis confirmed significant main effects of delay and decision, as well as a significant interaction (Figure S7, Table S2), indicating that neuronal delay coding is modulated by intertemporal choice behaviour.

### Impulsive individuals encode decisions just before reporting and show diminished prospective temporal coding

Impulsivity is characterized by a reduced ability to wait for future rewards and by a predisposition towards spontaneous and rash decisions. Although high impulsivity is linked to several psychiatric conditions, the neuronal mechanisms underlying impulsivity in the human brain are still widely unexplored. We hypothesise that impulsive individuals exhibit distinct neuronal decision coding and reduced coding for the temporal structure of rewards. To test this, we grouped participants into a high-impulsivity and a low-impulsivity group using a median split based on the individual discounting parameter *k*, and explored whether and how impulsivity affects the neuronal coding of upcoming decisions. Population-decoding analysis of neuronal activity sampled separately from participants with high vs. low impulsivity revealed distinct temporal profiles of the neuronal decision coding. In non-impulsive individuals, MTL neurons encoded decisions both earlier and more sustained than in impulsive individuals (Figure 4A). In contrast, for impulsive individuals, decisions were only decodable shortly before the end of the evaluation time interval, i.e., shortly before decisions were reported (Figure 4A). Interestingly, decision coding exceeded chance level already shortly before the onset of the choice options in non-impulsive individuals, indicating preparatory neural processes that were absent in the impulsive group (Figure 4A).

**Figure 4.**
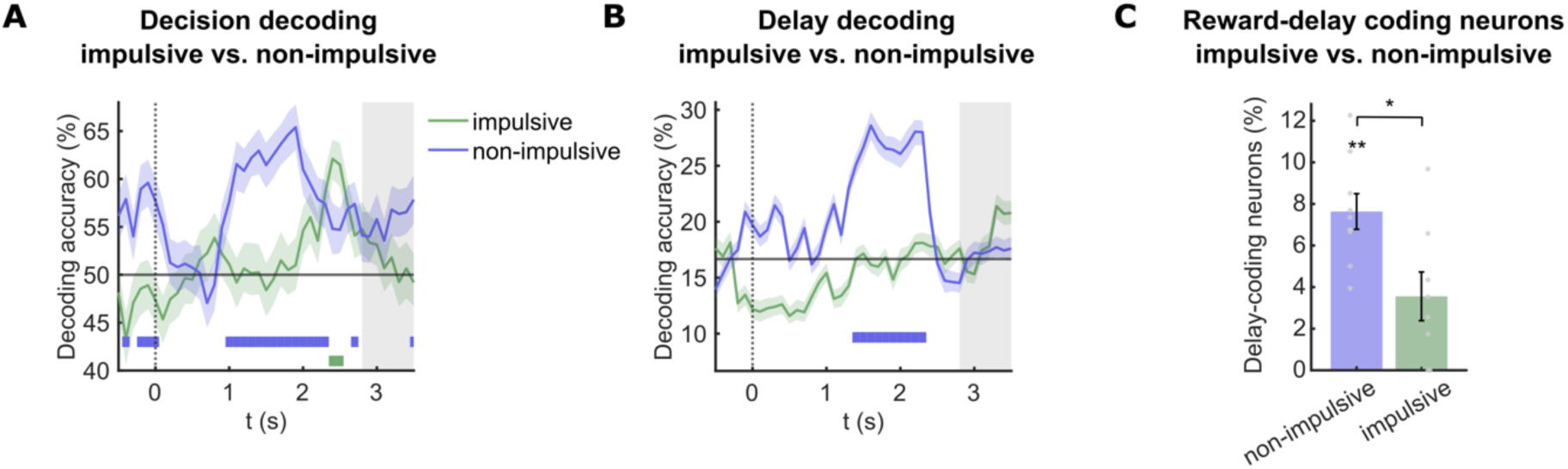
Impulsive individuals encode decisions just before reporting and show diminished prospective temporal coding. (A) Decision coding in human MTL neurons of impulsive and non-impulsive individuals (mean ± bootstrapped 95% CI). Participants were divided into two groups based on their individual discounting behaviour (median split of k values). Significant decision coding (horizontal thick bars, after FDR correction as in Figure 2E) was observed earlier and more sustained in non-impulsive individuals, whereas impulsive individuals’ decisions were decodable only shortly before the decision had to be reported. Non-impulsive individuals demonstrated significant decision coding already during the baseline interval before choice presentation. (B) Reward-delay coding in human MTL neurons of impulsive vs. non-impulsive individuals (mean ± bootstrapped 95% CI, median split as in A). Significant delay decoding was observed only in non-impulsive participants (thick horizontal blue bar). (C) Fraction of reward-delay coding neurons (mean ± s.e.m.) across individuals in the impulsive and non-impulsive groups. A significant fraction of reward-delay-coding neurons was observed in non-impulsive individuals, but not in impulsive individuals (non-impulsive: 7.6 ± 0.86%, P = 0.0078, N = 9; impulsive: 3.6 ± 1.2%, P = 0.9, N = 8, one-sided Wilcoxon signed-rank tests versus 5% chance), with significantly higher fractions found in the non-impulsive group (Impulsive vs. non-impulsive: P = 0.011, N = 8 vs. 9, two-sided Wilcoxon rank-sum test).

To further explore how neuronal representations of reward delays differ between impulsive and non-impulsive individuals, we trained decoders to predict reward delay based on neuronal populations sampled separately from the two groups. Notably, significant coding of reward delays was observed in non-impulsive individuals but was not detected in impulsive individuals (Figure 4B). Additionally, reward-delay coding neurons were significantly more prevalent in non-impulsive than in impulsive individuals (Figure 4C, P = 0.011, N = 8 vs. 9, two-sided Wilcoxon rank-sum test). These results suggest that high impulsivity is accompanied by impaired neuronal representations of the temporal dimension of future rewards. Training and evaluating decoders at the individual-subject level and comparing performance across impulsive and non-impulsive participant groups yielded consistent results (Figure S3). Comparisons of impulsive and non-impulsive individuals per anatomical region revealed significant delay and decision coding specifically in the hippocampus of non-impulsive individuals (Figure S4).

In conclusion, our results reveal distinct patterns of neuronal coding in the MTL of impulsive individuals, characterised by late-emerging decision coding and diminished representation of the temporal aspects of future rewards.

## Discussion

Although impulsivity in intertemporal decision-making governs many aspects of human behaviour, the underlying neuronal and circuit mechanisms of such decisions remain elusive. The behavioural effects and macroscopic brain dynamics underlying human delay discounting have been extensively studied ^7,39^. While non-invasive brain-imaging studies are informative at the systems and network level, they lack the spatial and temporal resolution to study physiological processes underlying delay and decision coding at the cellular level. Recordings of individual neurons in primates have significantly advanced our understanding of value and decision coding ^17^. Consequently, human single-neuron recordings present a rare opportunity to directly link human behaviour to neuronal activity, bridging animal and human research ^16,17^. Here, we recorded single-neuron activity in the human MTL during intertemporal choices to identify how decisions and temporal reward delays are encoded at the neuronal level. Our recordings revealed decision-coding neurons in the human amygdala and hippocampus. The population activity of MTL neurons predicted upcoming decisions, specifically in the hippocampus and amygdala. Intertemporal decisions require neuronal representation and processing of reward delays. We identified delay-coding neurons in the human hippocampus and entorhinal cortex that change their firing based on future reward delays. Decoding analyses revealed that MTL neurons, specifically hippocampal neurons, encode reward delays of the choice options. To explore neuronal correlates of impulsivity, we compared decision and delay coding between impulsive and non-impulsive individuals. In impulsive individuals, neuronal activity coded decisions only shortly before they reported their choice, whereas prolonged and anticipatory decision coding was found in non-impulsive individuals. Impulsive individuals additionally showed a diminished neuronal representation of future reward delays. Together, our findings demonstrate how individual MTL regions contribute to intertemporal decisions and reveal differences between impulsive and non-impulsive individuals at the neuronal level in humans.

### Prospective temporal coding by hippocampal and EC neurons

The human brain’s ability to perceive and represent time is fundamental to a variety of cognitive functions, including episodic memory and decision-making. Here, we discovered neurons in the human hippocampus and entorhinal cortex that encode future time intervals up to a year. Previous studies in rodents have identified neurons in the hippocampus that change their firing during brief delay periods in tasks analogous to delay discounting ^40^. Recordings in rodents also revealed hippocampal ‘time cells’ that represent temporal information, helping to organize experiences in time and supporting the hippocampus’s role in episodic memory and temporal sequencing ^41,30^. Temporal coding of task-relevant information was also observed in the hippocampal and EC neurons of primates ^42^, and more recently, time cells were identified in the human hippocampus and EC, where their activity was linked to episodic memory ^31,43^.

Previous research has focused on short, task-relevant time windows lasting seconds to hours ^30,41,43,44,31^, leaving the coding of longer prospective intervals, ranging from weeks to months, largely unexplored. Such extended planning and future thinking may represent a distinctively human capability ^45^. Our findings reach beyond earlier results by identifying coding schemes underlying long-term prospective temporal coding at the cellular level. Notably, we found delay-coding neurons in regions previously linked to temporal coding of past and present time. This suggests that common neuronal mechanisms may encode past, present, and future temporal information. Further research needs to elucidate the neuronal interplay of these temporal codes in humans and determine whether prospective temporal coding over extended periods and mental time travel are uniquely human features ^45^. Intertemporal choice requires integrating reward and delay information. The observed interaction between delay and decision coding suggests that human MTL neurons dynamically combine these variables during choice evaluation. Integrating eye-tracking data into future single-neuron studies of delay discounting will allow a finer-grained temporal analysis of the neuronal decision dynamics. In our task, we used fixed delays between 1 and 52 weeks but did not measure participants’ subjective perception of these intervals. Although humans can mentally simulate extended future periods ^35,46^, our behavioural and neuronal effects may reflect subjective delay weighting during intertemporal choice rather than a direct encoding of physical time. Future work should test how even longer past and future intervals are represented at the single-neuron level in humans, and how this relates to individual perception of time.

### Decision coding in the human hippocampus and amygdala

Whether and how the neuronal activity in the human MTL contributes to intertemporal decisions has been a long-standing question. Our findings reveal that hippocampal and amygdala neurons predict intertemporal decisions before they are reported.

In the rodent hippocampus, the activity of individual neurons predicts future route decisions through a prefrontal–thalamo–hippocampal circuit ^47^. Our results complement these findings by showing decision coding in the human hippocampus, suggesting that corresponding circuit coding may be important for human decision-making. Given the prefrontal cortex’s critical role in decision-making ^48^, coordinated neuronal interactions between the MTL and prefrontal cortex may be essential for supporting human intertemporal decision-making ^49,50^. Our finding of simultaneous decision and reward delay coding in the human hippocampus highlights its involvement in integrating temporal and choice information.

Human single-neuron recordings have provided evidence that amygdala neurons are important for coding decision-relevant variables including value ^51^, decision confidence ^52^, and personal valence ratings ^53^. Single-neuron recordings in nonhuman primates have shown that amygdala neurons predict economic decisions ^16,54^. Our finding of decision coding by human amygdala neurons provides converging evidence for the amygdala’s central role in decision-making. The observation that amygdala lesions impair human economic decisions ^12,13^ suggests that neuronal decision coding mechanisms in the amygdala shape human decision-making.

While the hippocampus’s role in memory processing and the amygdala’s role in affective processing are well established, their complex neuronal responses might contribute to higher-order cognitive functions through multidimensional coding ^55,56^. Our results extend response patterns in these regions to intertemporal decision-making, which involves integration of personal values, extended time periods, and future planning. Recording the same neurons across tasks may further elucidate how such complex information is integrated.

Converging evidence suggests a role of the hippocampus in decision-making through flexible integration of memory-related information into value-based decisions ^36^. Our findings of decision and delay coding by human hippocampal neurons provide support for this account, suggesting that hippocampal circuits integrate prospective simulations of value based on existing prior experience. The hippocampal and entorhinal formations contain spatially tuned neurons, including hippocampal place cells and entorhinal grid cells ^57,58^. More recent work suggests that this system also represents abstract relational information in cognitive-map-like formats ^59–61^. Primate studies further suggest that abstract value spaces are represented in the hippocampus and ventromedial prefrontal cortex ^62,63^. Our paradigm did not involve controlled trajectories through abstract value space, yet the observed representations of delay and decision information may be in line with navigating through cognitive maps of decision-relevant information. Although EC contained a significant fraction of delay-coding neurons, the absence of monotonically delay-correlated firing suggests a distinct coding principle compared to the Hp, potentially resembling grid-like schemes observed in spatial navigation and semantic maps ^64,59,61^. Future studies should explore whether the human EC encodes both temporal and spatial dimensions through a shared grid-like code.

### Impulsivity alters MTL decision and temporal coding

Impulsivity manifests early in life and has a fundamental impact, as the inability to delay gratification is linked to poorer social, economic, and academic outcomes ^65^. Additionally, impulsivity is a hallmark of multiple psychopathological conditions, including obsessive-compulsive disorder, substance abuse, and attention-deficit hyperactivity disorder ^4,5,66^. Strong delay discounting is a robust marker of impulsivity, consistently related to real-world behaviours such as substance abuse ^2,67,68^, financial mismanagement ^69,70^, and risk-taking tendencies ^71^. To better understand these conditions and behaviours, it is crucial to study the neuronal and circuit mechanisms underlying impulsive behaviour in humans.

We discovered that decisions in impulsive individuals are decodable only shortly before they are reported, while non-impulsive individuals exhibit earlier and more sustained decision coding. These results are in line with the idea that non-impulsive individuals plan upcoming decisions and reevaluate their decisions based on the choice options. Hippocampal lesions in rodents have been shown to bias animals toward more impulsive discounting behaviour ^72^, while hippocampal lesions in humans appear to affect discounting behaviour primarily when future episodic thinking is an integral part of the task^28,73^.

In non-impulsive individuals, we observed coding of the upcoming decisions even before the choice options were presented. As delays were pseudo-randomly organised and decision parameters were constantly updated, participants could not know the upcoming choice options. Decision coding before option presentation therefore likely reflects the intention and preplanning of future decisions. This decision-planning signal was only observed in non-impulsive individuals, indicating future decision planning as an integral part of non-impulsive behaviour. Interestingly, this signal was observed exclusively in the amygdala, suggesting that the human amygdala plays a role in intentional planning of upcoming decisions. Building on prior single-neuron recordings in nonhuman primates ^74^ and human imaging studies ^75^ that suggested the amygdala is key for planning upcoming economic decisions, our findings of decision coding before choice-option presentation provide single-neuron evidence for this involvement. While we used a median split to detect group-level differences in impulsivity, future studies with larger samples are needed to capture more fine-grained, continuous variations in decision and delay coding with impulsivity. For instance, applying non-invasive brain imaging in large cohorts may reveal how disruptions in delay and decision coding relate to clinical conditions such as substance abuse. Impulsivity is a multifaceted construct that includes not only impulsive decision-making but also failures of behavioural control, such as the inability to wait or suppress premature actions. While our study focuses on choice-related impulsivity measured through intertemporal choices, future research could investigate the extent to which different forms of impulsivity (e.g., impulsive actions and waiting impulsivity) share underlying neuronal mechanisms across brain networks including cortical-striatal and limbic networks ^76^.

We discovered that MTL neurons code for future reward delays specifically in non-impulsive participants, suggesting that insufficient neuronal representation of future times is part of the neuronal mechanisms underlying impulsivity ^7,29^. This effect may be moderated by a reduced capability for future episodic thinking, which could be facilitated by hippocampal prospective temporal coding ^27–29^.

Collectively, we demonstrate the involvement of the human MTL in intertemporal decision-making, revealing decision coding by amygdala and hippocampal neurons, and prospective temporal coding in hippocampal and entorhinal neurons. Importantly, we show how impulsivity influences both decision and temporal coding in the MTL, paving the way for future research on impulsivity in health and disease.

## Methods

### Delay discounting paradigm

Participants were asked to choose between a larger-but-later (LL) amount of 10 € at a variable delay *D* (1, 2, 5, 11, 24 or 52 weeks) and a smaller-but-sooner (SS) immediate reward. Delays were presented in pseudorandom order evenly distributed across the entire experiment. Participants made 15 choices for each delay while the SS amount was adjusted based on previous responses to account for individual differences in the discounting behaviour. The SS reward was calculated as:

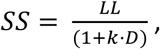

with *k* denoting the discounting parameter ^77^. In the first trial, the discounting parameter was set to *k* = 0.02 for all participants ^77,78^. Choosing SS over LL indicates a lower subjective value of LL. Consequently, SS was decreased by increasing *k* ^38^. If LL was chosen, *k* was decreased for a higher SS. To allow fast convergence towards the individual discounting parameter in the beginning (first 20 trials), *k* was increased or decreased by 10% of its previous value (*k_t+1_* = *k_t_* ± *k_t_·0.1*; with *t* denoting the trial number). To allow a more fine-grained variation in the remaining trials, *k* was adjusted by 5% of its previous value thereafter (*k_t+1_* = *k_t_* ± *k_t_·0.05*). Participants were divided into high- and low-impulsivity groups through a median split based on their individual discounting parameter (high impulsivity: *k* > 0.021, N = 8; low impulsivity: k ≤ 0.021, N = 9). Participants were instructed to consider each of their choice options carefully. As prior research has demonstrated consistent discounting for real and hypothetical rewards ^79–81^, we refrained from actual monetary payouts in our clinical setting.

Each trial started with a fixation cross (1 s ± 200 ms) before choice options were presented simultaneously above or below the centre of the screen in pseudorandom order. Monetary rewards were presented next to the delay information. Participants were instructed to evaluate choice options thoroughly and to respond with a button press once the central fixation point turned red (3 s ± 200 ms after choice-option onset). The experimental protocol was implemented in MATLAB 2019a (MathWorks, Natick, MA) using Psychtoolbox3 ^82–84^.

### Subjects

We recorded single-neuron activity in 17 pharmaco-resistant epilepsy patients (12 female, 5 male; aged 37.6 ± 12.1 years (mean ± s.d.)) undergoing invasive pre-surgical seizure monitoring. Data were recorded at the Department of Epileptology at the University of Bonn Medical Center, Bonn, Germany. This study adhered to and was approved by the Medical Institutional Review Board of the University of Bonn, Germany (license no. *147/19*). Informed written consent was provided by every patient.

### Human Single-Neuron Recordings

Single-unit activity was recorded from bilaterally implanted Behnke-Fried depth electrodes in the amygdala, hippocampus, entorhinal cortex and parahippocampal cortex. Target locations were verified based on post-implantation CT scans co-registered to pre-implantation structural MRI scans using the LeGUI toolbox ^85^. Recordings from electrodes outside MTL target regions were not analysed. Microwire bundles were manually trimmed and inserted through the clinical electrodes such that they protruded 3-5 mm from the tip of the clinical electrodes. Each platinum-iridium microwire bundle contained eight high-impedance recording microwires and one low-impedance reference wire (Ad-Tech Medical, Racine, WI). Microwire signals were recorded with an ATLAS system (Neuralynx, Bozeman, MT) using a sampling rate of 32,768 Hz and a bandpass filter of 0.1 to 9,000 Hz. Spike extraction and sorting were performed with Combinato ^86^. Only spikes of negative polarity were extracted. Spike shapes shown in this manuscript have been inverted for visualisation purposes. We recorded neuronal spiking activity from 1,002 units (461 single units and 541 multiunits). Spike-sorting quality was manually verified based on spike shapes, signal-to-noise ratios, inter-spike-intervals and cross-correlations, and artefacts were manually rejected. No statistical methods were used to predetermine sample sizes, but the participant and neuronal sample sizes are within the range of recent human single-neuron studies ^87,88^.

### Statistical Analyses

#### Individual Discounting Behaviour

Participants’ individual discounting parameters *k* were estimated by fitting a logistic growth curve based on individuals’ behaviour. The function

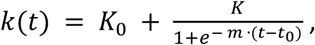

was fitted to the individual course of adjusted *k* parameters using MATLAB’s ‘*fit.m’* function. Here, *t* is the trial number and *K*_0_, *K*, *m*, *t*_0_ are the resulting parameters of the fit (start values (*K*_0_, *K*, *m*, *t*_0_) = (0, 0.02, 0.1, 45) and fit range of (*K*_0_, *K*, *m*, *t*_0_) = ([0, *inf*], [0, 10], [−0.5, 0.5], [−90, 90])). Individual discounting parameters for each participant *i* were estimated as the limit value for an infinite number of trials: 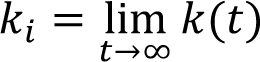.

### Decision- and Delay-Coding Neurons

To identify decision-coding neurons, we performed a two-sided t-test comparing the trial-wise spike counts between trials in which the SS option was chosen and trials in which the LL option was chosen (P < 0.05). Spike counts for each trial were calculated during choice evaluation using a 200 to 2,800 ms time window after choice option onset. Similarly, reward delay-coding neurons were identified based on a one-way ANOVA (P < 0.05) of the trial-wise spike counts across reward delays using the same time window. We further identified delay-correlated neurons by testing whether each neuron’s spike count was significantly correlated with delay duration using a Spearman correlation. To assess delay and decision coding as well as their interaction on the population level, we used generalized linear mixed-effects models (GLME) to predict trial-wise spike counts (SC) of each neuron during the delay evaluation period (200 to 2,800 ms). Decision (SS or LL), delay rank (1 to 6) and their interaction were included as fixed effects, with a random intercept for each neuron. Poisson models were fitted using the restricted maximum pseudo-likelihood estimation with a log link function implemented in MATLAB’s ‘fitglme’ function.

### Neuronal Decoding Analysis

Neuronal decoding analyses were employed to investigate how trial-wise decisions and delays are encoded by the activity of MTL neurons. Multi-class maximum-correlation classifiers implemented in the Neural Decoding Toolbox ^34^ were used for all decoding analyses. Decoding was based on the pseudo-population of recorded neurons across recording sessions without any prior selection of significantly decision- or delay-coding neurons. For each anatomical target region, decoders were trained based on all recorded neurons using 10 cross-validation data splits and 100 resampling runs. To enable this 10-fold cross-validation, only sessions in which both choice options were selected at least 10 times were included in the decision decoding analysis (N = 14). Spike counts of each neuron were z-normalized across trials before decoding. Neuronal spiking activity was binned using a 1 s moving time window and 100 ms step size, covering the interval from -1 s to 4 s after choice option onset (i.e., 41 time-bins). To investigate the impact of the moving time-window length on the decoding results, we repeated the decision and delay decoding analysis using various time windows ranging between 100 ms and 3,000 ms with steps of 100 ms. The results demonstrate consistent decision and delay coding across various time windows (Figure S5). To assess how each delay option contributed to delay coding, we used the diagonal elements of the 6-class classifier’s confusion matrix to quantify decoding accuracy for each delay (Figure S6).

Decoding results are shown as the mean decoding performance across resampling runs. Decoding variability is visualised as 95% bootstrapped confidence intervals across resampling runs, computed based on MATLAB’s ‘bootci’ function using 1,000 bootstrap samples. To estimate statistical significance, the decoding was repeated 1,000 times on label-permuted data using identical classifiers. The decoding performance of these label-permuted data was used to estimate a surrogate distribution across all time bins. For each time bin, P values were estimated based on the percentile of the real decoding results in the surrogate distribution of all label-permuted decoding results (P = 1 - percentile). False-discovery-rate (FDR) correction was applied to correct for multiple testing across time bins (N = 41).

## Data Availability

The data supporting the key findings of this study and needed to reproduce the main figures in this manuscript will be made publicly available upon publication.

## Acknowledgements

We thank all the patients for their participation in this research. This research was supported by grants from the DFG (SFB 1436 A03 (SD), MO 930/4-2, MO 930/15-1, SPP 2411), BMBF (031L0197B) (F.M.) and the NRW Network Grant (iBehave) (F.M.). The funders had no role in study design, data collection and analysis, decision to publish or preparation of the manuscript. The icons included in the figures were obtained via a Noun Project NounPro subscription (https://thenounproject.com).

## Author Contributions

S.D. conceived and designed the experiment. S.D. and M.K. implemented the experiment. F.M. and R.S. recruited patients. V.B. and F.M. implanted the electrodes. M.K. collected and curated all data. M.K. and S.D. analysed the data. M.K., S.D. and F.M. interpreted the data and wrote the manuscript. F.M. administered the project. All authors discussed the results and commented on the manuscript.

## Supplemental Information

**Table S1.**
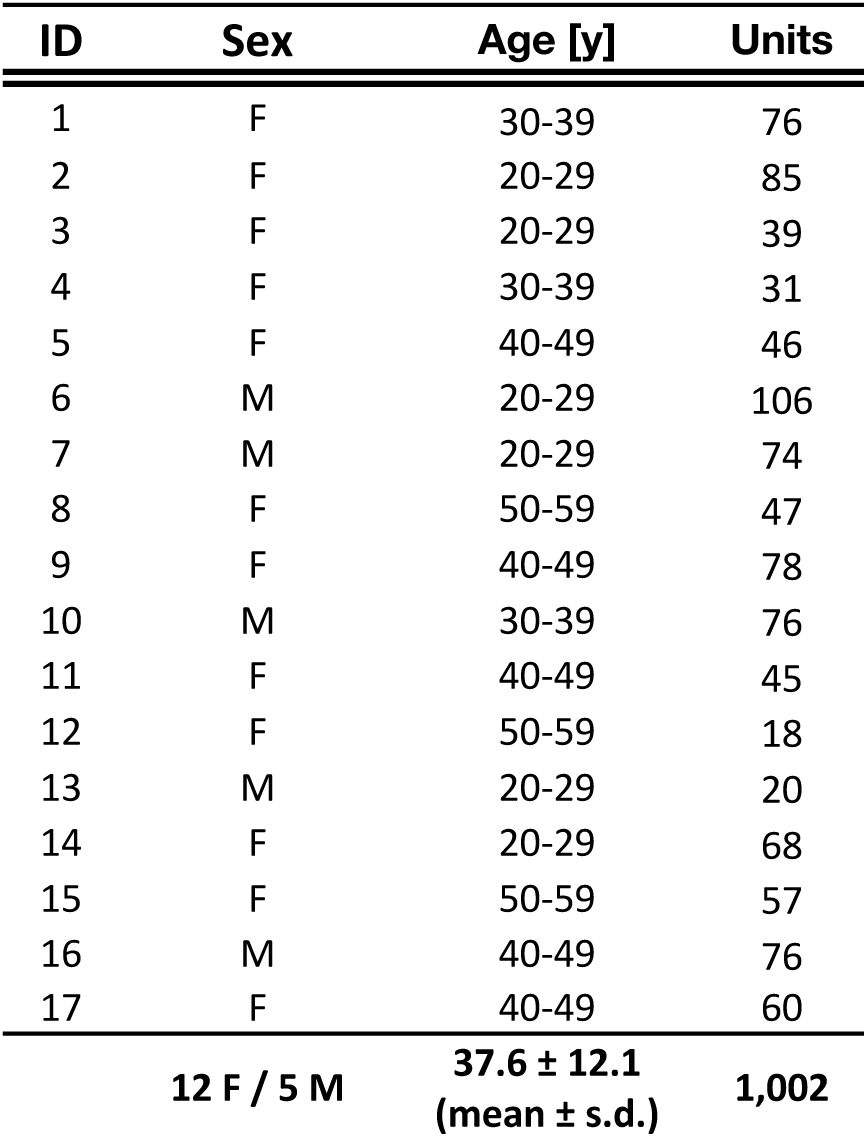
Overview of recordings and patients. Participant characteristics, including age range, sex and number of recorded units per participant (rows). The final row shows the total number of units and the mean age of all 17 patients. Patient IDs are arbitrarily assigned from 1 to 17. Age is reported in ranges to protect patient confidentiality.

**Table S2.** Generalized linear mixed-effects model. General linear-mixed effects model (GLMEs) predicting trial-wise spike counts (SC) of neurons (N = 1,002) during delay evaluation (200 – 2,800 ms) based on decision (SS chosen), delay and their interaction. Significant main effects of decision and delay, as well as a significant decision-by-delay interaction, were observed. Model: SC ∼ 1 + Decision x Delay + (1 | NeuronID). A random intercept was included for each neuron. Poisson models were fitted using a restricted maximum pseudo-likelihood estimation with a log link function.

| Fixed Effect | Estimate | Std. Error | t-value | P-value | 95%-CI |  |
| --- | --- | --- | --- | --- | --- | --- |
| **Intercept** | 1.022 | 0.039 | 25.93 | $< 10^{-10}$ | [0.945 | 1.099] |
| **Decision** | -0.016 | 0.007 | -2.24 | 0.02 | [-0.031 | -0.002] |
| **Delay** | -0.005 | 0.001 | -3.84 | 0.0001 | [-0.007 | -0.002] |
| **Decision x Delay** | 0.0042 | 0.0019 | 2.25 | 0.02 | [0.0005 | 0.0078] |

**Figure S1.**
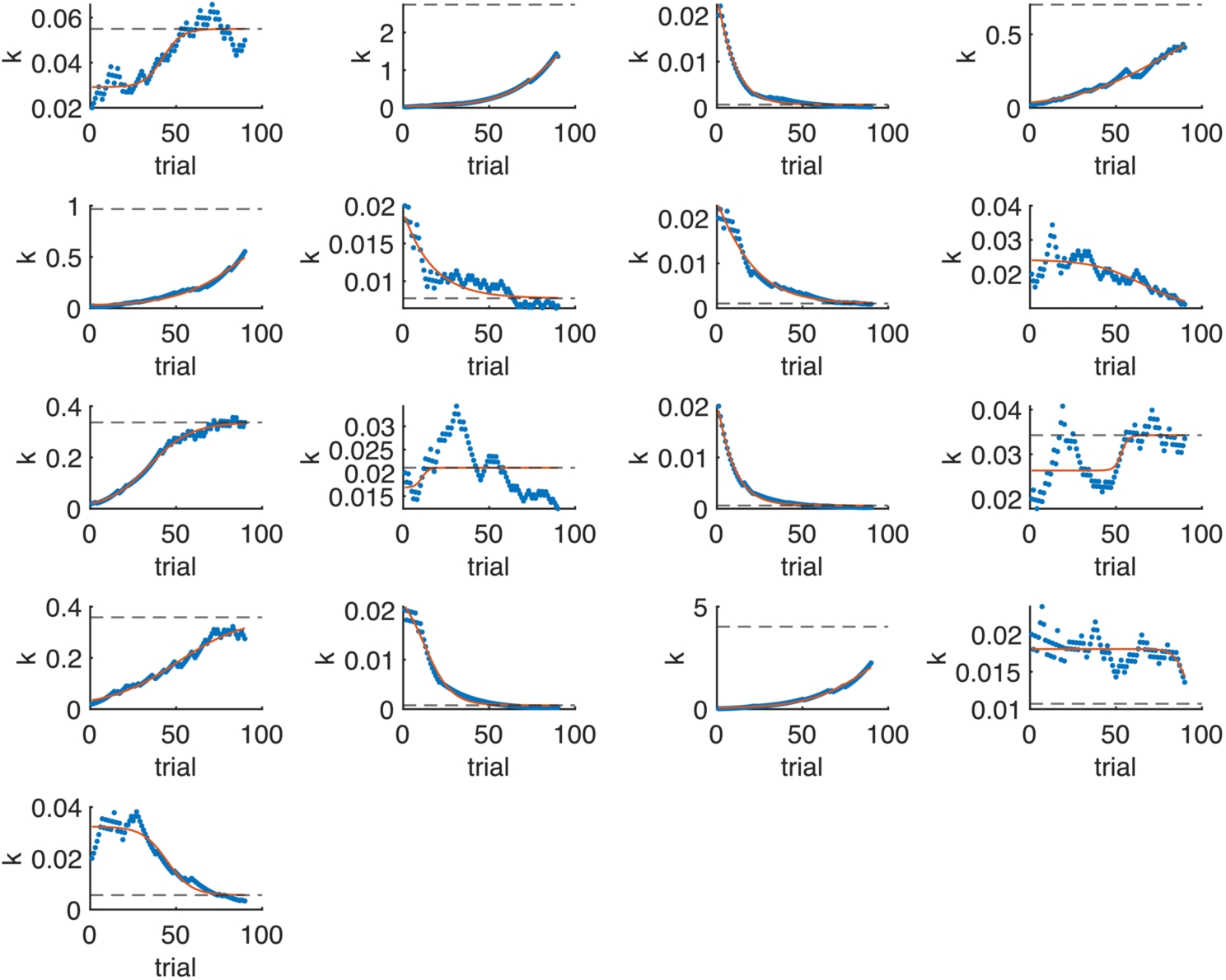
Behaviour per participant. Each panel illustrates the adaptive evolution of the k-parameter for an individual participant across all 90 decisions. The red line represents the logistic growth function fitted to the data, with the dashed horizontal line indicating the estimated k-parameter from this fit.

**Figure S2.**
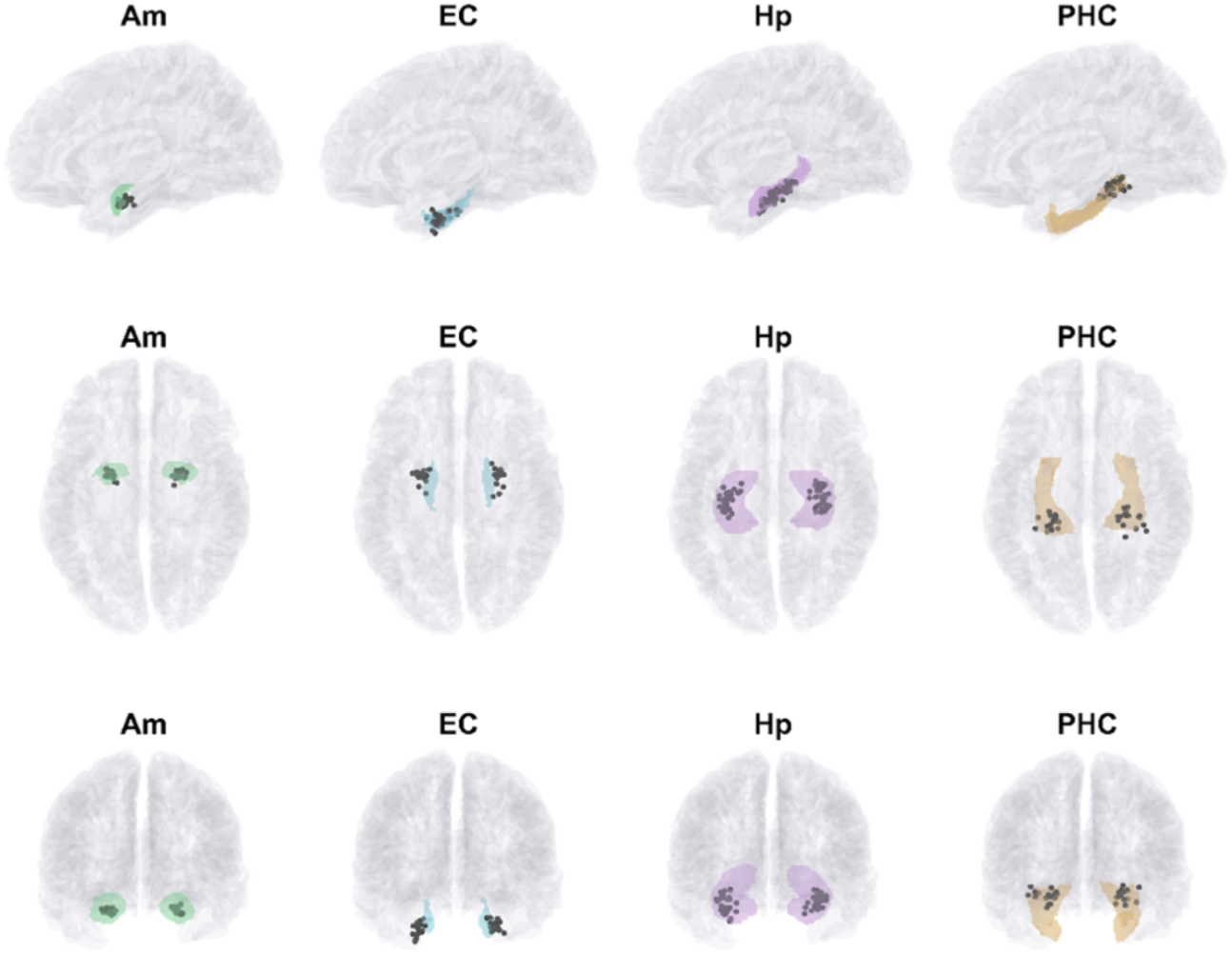
Electrode locations across anatomical target regions and participants. Black dots indicate innermost clinical electrodes projected to the MNI-ICBM152 template. Electrode locations are shown in sagittal, axial, and coronal view (top to bottom), with electrodes projected to a single hemisphere in the sagittal view. Electrode locations were visualised using FieldTrip ^89^ and the custom MATLAB function ‘plot_ecog’ (https://github.com/s-michelmann/moment-by-moment-tracking/blob/master/plot_ecog.m). Anatomical target regions are highlighted for visualization using the connectivity-based parcellation of the Brainnetome Atlas ^33^. Note that all surgical planning and post hoc electrode localization were performed in each patient’s native space.

**Figure S3.**
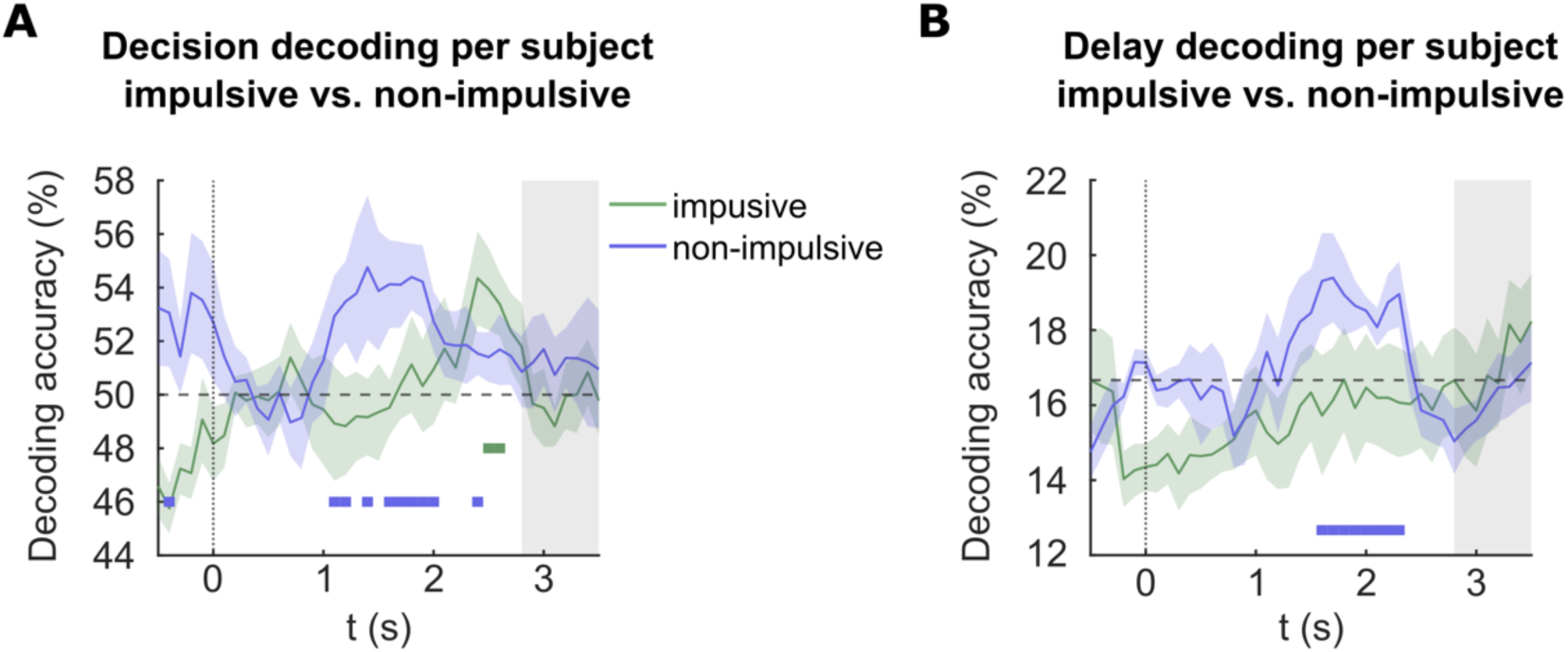
Delay and decision decoding per participant. (A) : Decision decoding accuracy (mean ± s.e.m.) across participant for impulsive (green) and non-impulsive (blue) participants. Decoding performed as in the main decoding analysis (Figure 4A) but separately for each participant based all recorded neurons. Thick horizontal bars indicate time points of significant decoding (one-sided Wilcoxon signed-rank tests against chance level across participants, uncorrected). Significant decision coding was observed for both impulsive and non-impulsive participants, but with earlier and more sustained decision coding in the non-impulsive group. Preparatory decision coding prior to option onset was only present in the non-impulsive group. (B) Delay decoding accuracy (mean ± s.e.m.) across participants for impulsive (green) and non-impulsive participants (blue). Decoding per participant as in (A). Thick horizontal bars below chance level (dashed line at 16.67%) indicate time points with significant above-chance decoding (P<0.05, one-sided Wilcoxon signed-rank tests against chance level across participants, uncorrected). Significant delay coding was observed only in the non-impulsive group.

**Figure S4.**
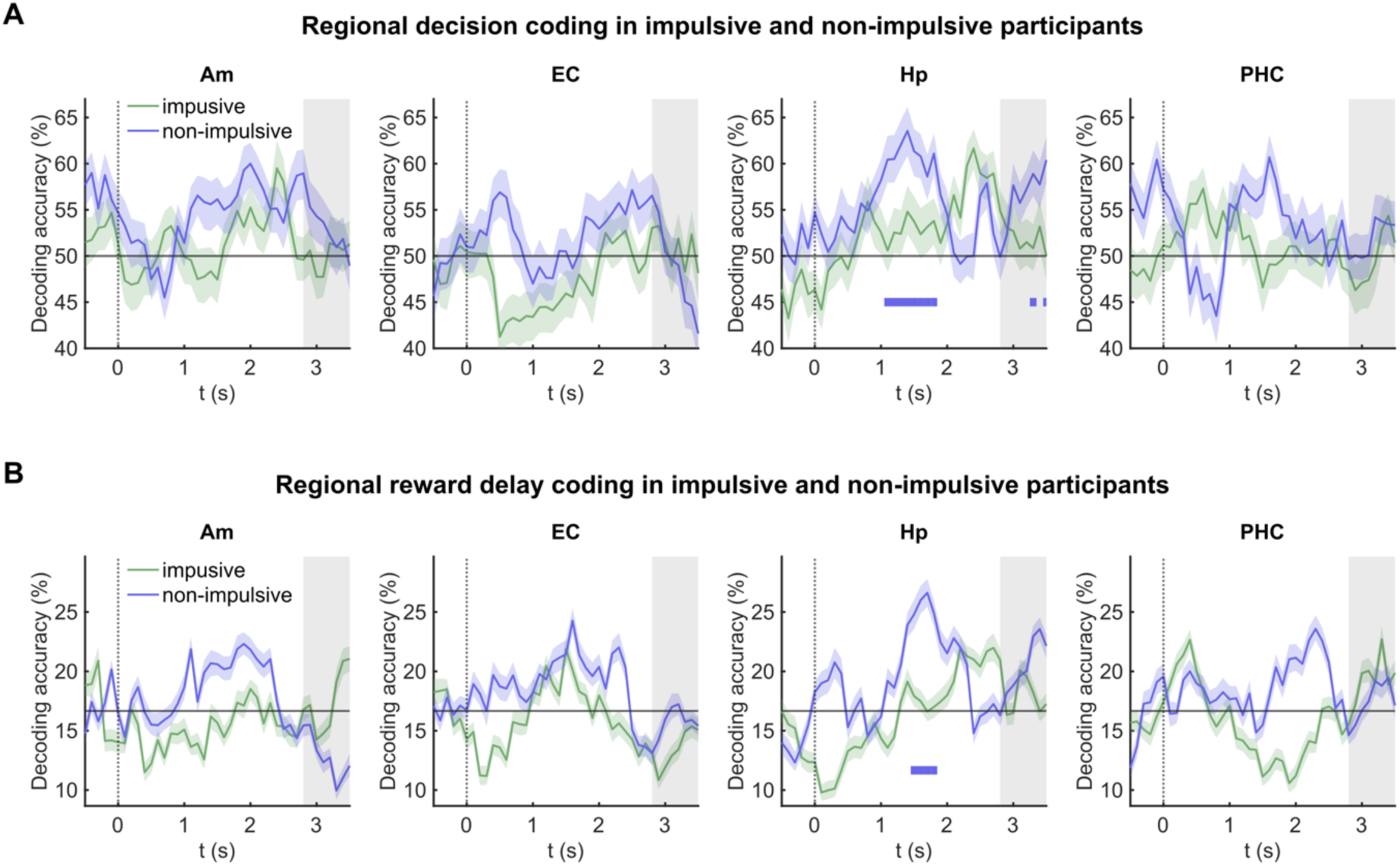
Regional decision and delay decoding in impulsive versus non-impulsive participants. (A) Decision decoding performance (mean ± bootstrapped 95% CI across resampling runs) by anatomical target region for the group of impulsive (green) and non-impulsive (blue) individuals (median split). Decoding and display as in Figure 4A, with decoding analysis performed separately for neurons per anatomical target region. Significant decision coding (horizontal thick bars, after FDR correction) was observed only in the hippocampus of non-impulsive individuals while it did not reach significance in the group of impulsive individuals. (B) Temporal delay decoding performance (mean ± bootstrapped 95% CI across resampling runs) by anatomical target region for the group of impulsive (green) and non-impulsive (blue) individuals (median split). Decoding and display as in Figure 4B, with decoding analysis performed separately for neurons per anatomical target region. Specifically, hippocampal neurons in non-impulsive individuals exhibited significant delay coding (horizontal thick bars, after FDR correction). Note, that subdividing the neuronal population by region and into an impulsive and non-impulsive group reduces the power of population decoding.

**Figure S5.**
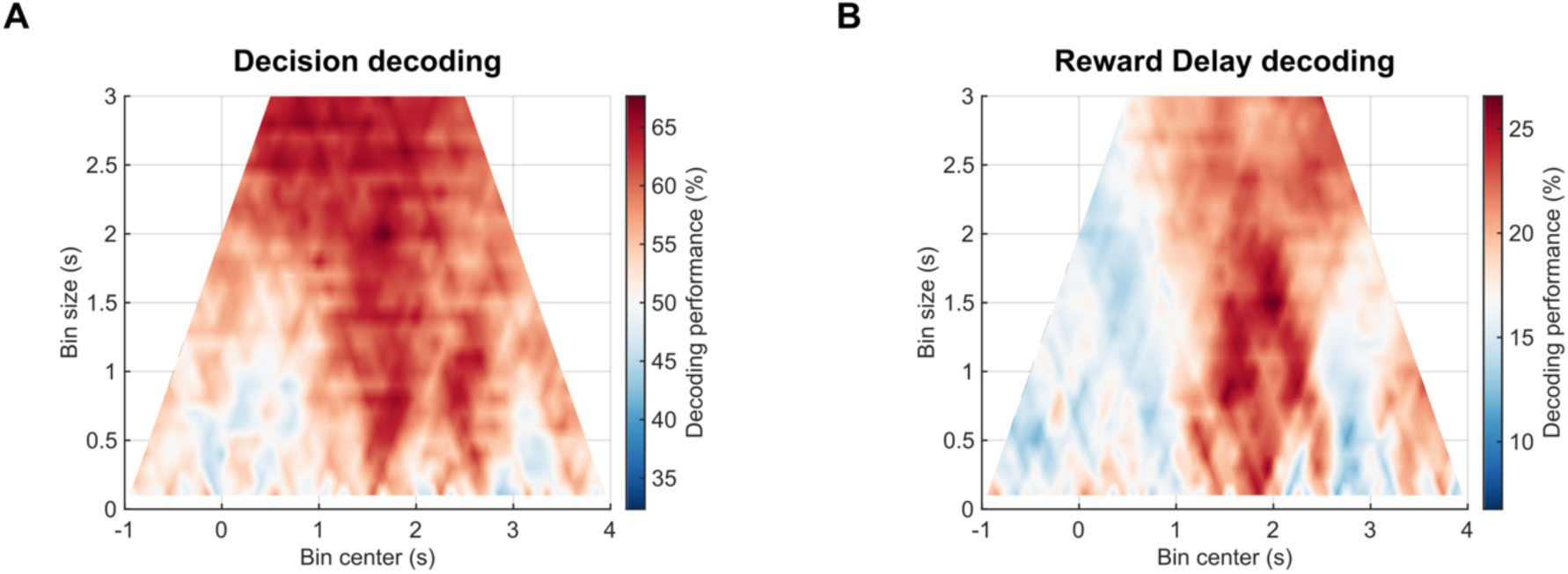
Decision and delay coding across decoding time windows. (A) Colours indicate decision decoding performance for different decoding windows (y-axis, bin sizes from 0.1 to 3 s in 100 ms steps) and time points relative to option onset (0, x-axis). Above-chance decoding performance was observed across a wide range of decoding time windows. (B) Delay decoding performance (colour-coded) across different decoding windows (y-axis, bin sizes from 0.1 to 3 s in 100 ms steps) and time points relative to option onset (0, x-axis). Delay information could be decoded across a wide range of decoding time windows during choice evaluation. All decoding analyses in this manuscript used a 1-second decoding window.

**Figure S6.**
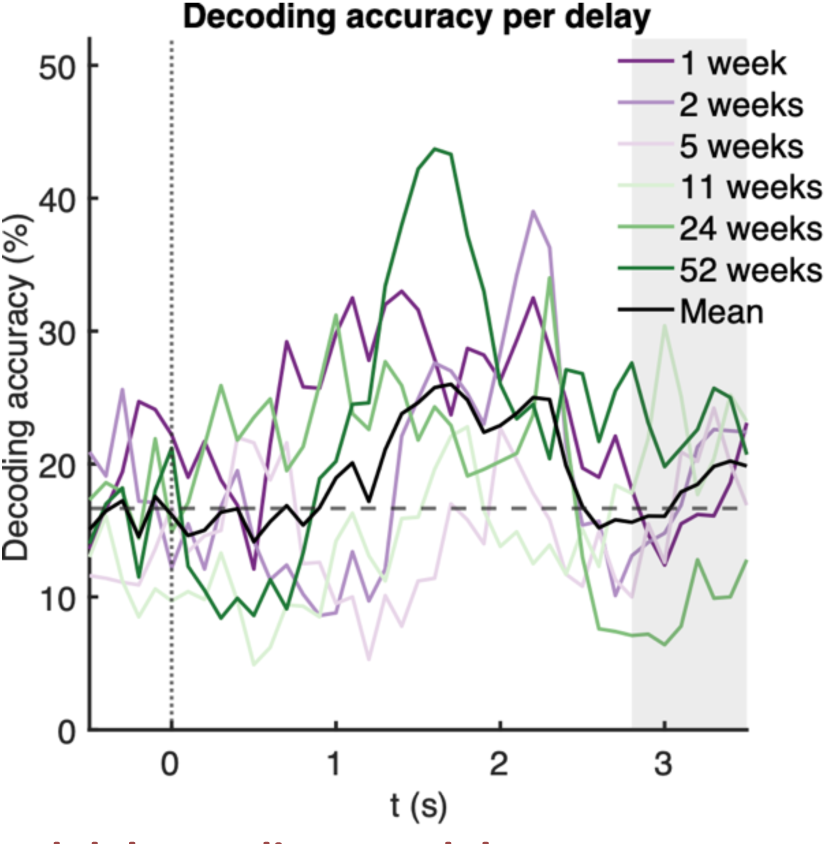
Prospective reward delay coding per delay. Reward-delay coding in human MTL neurons as a function of time (as in Figure 3E). Coloured lines represent classification accuracy for each delay period, with the black line indicating the overall performance of the 6-class classifier. The diagonal elements of the 6-class delay classifier’s confusion matrix were used to evaluate classification performance for each individual delay. Chance level performance (16.67%, i.e., 1 out of 6 delays) indicated by the dashed horizontal line. Note that the shortest (dark purple) and longest delays (dark green) show the highest decoding accuracy, suggesting enhanced neural discriminability at the temporal extremes.

**Figure S7.**
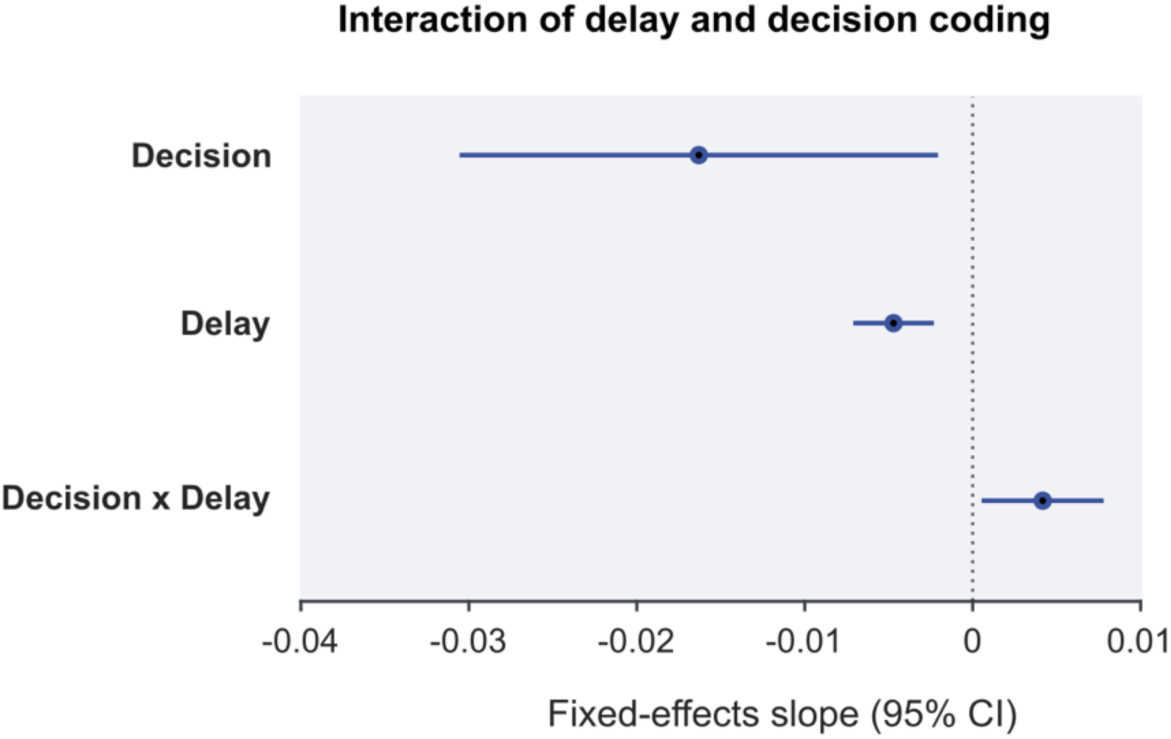
Interaction of delay and decision coding in the human MTL. Forest plot of the estimated fixed-effects slopes and 95% confidence intervals for the main effects of delay, decision, and their interaction on neuronal spiking in the GLMM fitted to all 1,002 human MTL neurons (Table S2). Both delay and decision exhibited significant main effects, and their interaction was also significant.

